# Implementing and assessing an alchemical method for calculating protein-protein binding free energy

**DOI:** 10.1101/2020.10.02.324442

**Authors:** Dharmeshkumar Patel, Jagdish Suresh Patel, F. Marty Ytreberg

## Abstract

Protein-protein binding is fundamental to most biological processes. It is important to be able to use computation to accurately estimate the change in protein-protein binding free energy due to mutations in order to answer biological questions that would be experimentally challenging, laborious or time consuming. Although non-rigorous free energy methods are faster, rigorous alchemical molecular dynamics-based methods are considerably more accurate and are becoming more feasible with the advancement of computer hardware and molecular simulation software. Even with sufficient computational resources, there are still major challenges to using alchemical free energy methods for protein-protein complexes, such as generating hybrid structures and topologies, maintaining a neutral net charge of the system when there is a charge-changing mutation, and setting up the simulation. In the current study, we have used the *pmx* package to generate hybrid structures and topologies, and a double-system/single-box approach to maintain the net charge of the system. To test the approach, we predicted relative binding affinities for two protein-protein complexes using a non-equilibrium alchemical method based on the Crooks fluctuation theorem and compared the results with experimental values. The method correctly identified stabilizing from destabilizing mutations for a small protein-protein complex, but was not as successful to the larger, more challenging antibody complex. In addition, the correlation between predicted and experimental relative binding affinities was high for smaller complex, and low for the other larger complex.

## INTRODUCTION

Protein-protein binding is an essential phenomenon in molecular biology, and directly mediates most functions in cells such as cellular metabolism, signal transduction, and coagulation among many other biological processes.^1,2^ Mutations of the amino acids in protein-protein complexes can modulate, or even disrupt protein-protein interactions by changing the associated binding free energy (Δ*G*) of the protein–protein complexes. The binding free energy of the protein-protein complexes determines the stability of association and the conditions for protein-protein complex formation.^3^ It is important to be able to quantify the stabilities of protein complexes and how they can be modified by amino acid mutations and how they are affected by evolution.

Many techniques have been employed to determine the change in the protein-protein binding free energy due to a mutation (i.e., relative binding affinity, ΔΔ*G*). Experimental biophysical and biochemical methods are routinely used, but these methods are laborious, expensive, time consuming, and are limited by technical challenges.^4–7^ By contrast, computational methods can be relatively inexpensive, and the accuracy of such methods has been improved with the advancement of computational resources and better forcefields.^8–10^ Computational methods for estimating ΔΔ*G* values can be broadly classified as either non-rigorous or rigorous.^11^

Non-rigorous free energy methods typically use a single, static all-atom structure of the protein complex. These methods typically have energy functions that are trained using experimentally measured binding affinities or changes in affinities.^12,13^ Many such semi-empirical approaches have been developed that combine molecular mechanics and various optimized energy terms from available experimental data.^14^ For example, BeAtMuSiC and mCSM use coarse-grained statistical potentials derived from known 3-D structures of proteins and machine learning.^15,16^ FoldX uses empirical forcefield trained by experimentally measured binding free energies or changes in affinities.^12,13^ Other so-called docking/scoring algorithms can predict binding affinities based on predicted binding poses and putative binding interactions between protein-protein complexes.^17–19^

Rigorous free energy approaches are based on the principles of statistical mechanics and use molecular simulations to explore the conformational space.^20^ These methods typically provide more accurate ΔΔ*G* predictions, compared to non-rigorous. One reason for this is that they inherently consider the conformational flexibility of the proteins and hence the entropic contribution. In recent years, rigorous approaches have made tremendous efficiency and theoretical advancements.^11,20^ Rigorous free energy calculation approaches are typically classified into three categories: endpoint methods, physical path sampling, and alchemical transformation.^20^ Endpoint methods typically use molecular mechanics force fields with implicit solvent models such as molecular mechanics generalized Born surface area (MMGB/SA) and molecular mechanics Poisson–Boltzmann surface area (MMPB/SA).^21,22^ These methods are computationally less expensive than other rigorous approaches since simulations are only performed for two states, however their accuracy is system-dependent and sensitive to simulation protocols such as sampling strategy and entropy calculation. For path sampling approaches, the physical unbinding and/or binding pathway of the protein with respect to its partner is sampled to obtain underlying the free energy profile connecting bound and unbound states.^23–25^ This category of methods can be very accurate but requires exhaustive conformational sampling along the pathway making it computationally expensive. Finally, alchemical methods exploit unphysical pathways by morphing, creating and annihilating atoms.^26–29^ These methods use molecular mechanics force fields as an energy function and the sampling of the correct thermodynamic ensemble is maintained by thermostatted and baroststed dynamics. The primary advantage is that the alchemical pathway does not need to be correlated with the physical binding process. This is particularly advantageous when considering relative binding affinity calculations due to single amino acid mutations (such as the current study). In this case, one need only calculate the free energy change due to alchemically mutating the amino acid to another type in both the bound and unbound states.

Rigorous molecular dynamics (MD)-based alchemical free energy calculation can be performed by using equilibrium (e.g., free energy perturbation,^30^ thermodynamics integration^31^), or nonequilibrium (e.g., the Jarzynski equality,^32,33^ Crooks fluctuation theorem^34^) methods. The initial simulation set up is the same for both equilibrium and non-equilibrium methods, but the protocols used during the simulations and post analyses are different. The Hamiltonian *H* is coupled to a parameter *λ* that navigates the system from wildtype (*λ* = 0) to mutant (*λ* = 1). While such alchemical methods can be very accurate, they can also be computationally expensive since sufficient sampling is required to overcome the energetic and entropic barriers. In addition, the initial set up is not user friendly, particularly when there is a change in the net charge of the system.^29,35,36^ Specifically, the set up requires the topology of the protein system to ensure all bonded and non-bonded interactions are correctly switched from *λ* = 0 to 1.

To enable more user-friendly alchemical free energy calculations, de Groot *et al*. developed a package called *pmx* that automatically generates hybrid protein structures and topologies using forcefield-specific pre-generated mutation libraries.^37–39^ Moreover, to maintain the net charge of the system during alchemical transformation, they developed an approach that uses two protein systems in a single simulation box (double-system/single-box).^37,40^ Their approach of using *pmx*-generated topologies with a double-system/single-box was previously used to predict protein folding ΔΔ*G* values due to mutations.^37,38^ Prior to the development of the *pmx package*, de Groot *et al*. used the hybrid topologies approach to calculate binding free energies for ubiquitin in complex with different protein substrates using a fast-growth thermodynamic integration approach with the Crooks Gaussian intersection (CGI) method.^41^ The main purpose of their study was to analyze ubiquitin conformations due to point mutations and predict the sign of ΔΔ*G* for binding different substrates. They studied 11 mutations and obtained a Pearson correlation coefficient of 0.70 (p = 0.016). However, they have not explored the transition time per snapshot for nonequilibrium simulations. Later the same group tested *pmx* with double-system/single-box approach to predict ΔΔ*G* binding free energies for the protein-protein complex of α-Chymotrypsin with its inhibitor Turkey Ovomucoid third domain with nine observed mutations of site L18 of Turkey Ovomucoid third domain.^40^ The correlation coefficient between predicted and experimental ΔΔ*G* was 0.80. Although promising, this protein-protein complex is small, all nine mutations occurred at the same amino acid site, and were non-charge mutations.

Here, we tested the performance of using *pmx* with a double-system/single-box approach in a systematic manner using two protein-protein complexes of different size with a wide range of experimental ΔΔ*G* values. For each system, we selected eight mutations from different sites with a broad range of experimental ΔΔ*G* values. We estimated ΔΔ*G* values using *pmx* hybrid topologies with a double-system/single-box approach, and the non-equilibrium CGI method. Predicted ΔΔ*G* values were compared with experimental values. Higher correlation was found for the smaller protein-protein complex compared to the larger, more complex, antigen-antibody system. Our results suggest that there is still room for improvement in rigorous binding free energy methods, especially for large, complex protein-protein systems.

## METHODS

### Test System Selection

We selected two protein-protein complexes from the SKEMPI database^42^ as test systems for this study. We chose the relatively small Barnase (110 aa) – Barstar (89 aa) complex (Protein Data Bank (PDB) ID: 1BRS)^43^ and the larger, more challenging, antigen–antibody complex of lysozyme (129 aa) – HY/HEL-10 FAB (429 aa) (PDB ID: 3HFM)^44^. Eight mutations were selected from the list of experimental ΔΔ*G* values for each system from the SKEMPI database. We chose these systems and mutations based on several criteria: (i) ΔΔ*G* values should vary in sign—important since mutations with negative (stabilizing) values are often more difficult to predict compared to positive (destabilizing) values; (ii) there should be a small number of missing residues in the 3-D structure of the protein complexes; (iii) chosen mutations should be non-alanine-scanning point mutations at differing amino acid sites; (iv) reported mutations should be on multiple chains (Figure 1).

**Figure 1.**
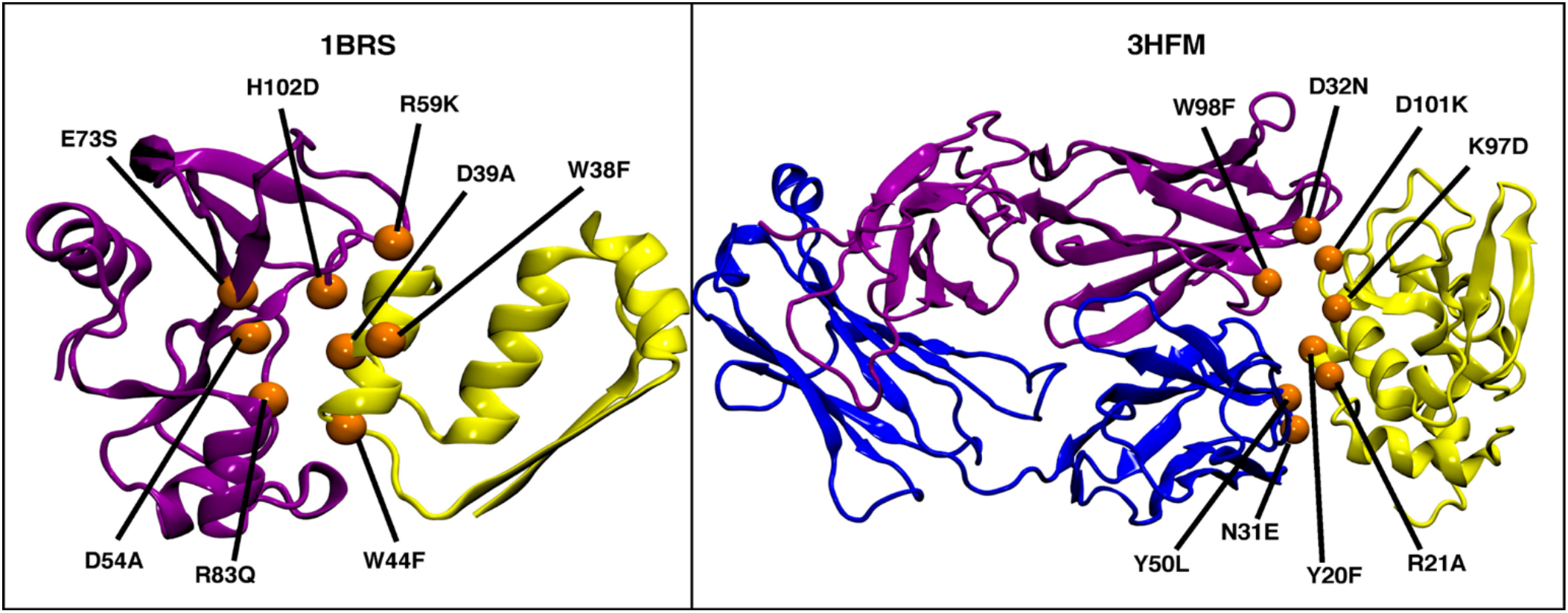
3-D structures of the test systems used in the current study with the eight selected mutations shown as orange spheres. Left: Barnase (purple) – Barstar (yellow) protein complex (PDB ID: 1BRS); Right: Lysozyme – HY (yellow) HEL-10 FAB (purple and blue) antigen– antibody complex (PDB ID: 3HFM).

### Preparation of Protein-Protein Complexes

The 3-D structures of protein-protein complexes were downloaded from the PDB server (https://www.rcsb.org) and edited to preserve only the coordinates of the two or three interacting chains listed in the SKEMPI database.^42^ All missing residues and atoms were then added using MODELLER software.^45^ Mutants were generated using the BuildModel command from FoldX software.^12,13^ This process provided nine input structures for each protein complex (a wild-type and eight mutant forms) to carry out alchemical free energy calculations.

### Construction of Hybrid Residues

Alchemical binding free energy calculations require the construction of a non-physical pathway of intermediate states connecting the wild-type amino acid (λ = 0) to its mutant form (λ = 1). The *pmx* webserver^37,38^ allows automatic generation of these intermediate states by producing hybrid amino acid states representing a mixture of wild-type and mutant form. (see Figure 2) Both wildtype and mutant complex structure files were uploaded to the *pmx* webserver. The *pdb2gmx* option to add hydrogen atoms, and the Amber99SB*ILDN modified force field options were selected. The *pmx* webserver output consisted of hybrid structure and topology files compatible with GROMACS to perform the alchemical molecular dynamics (MD) simulations.

**Figure 2.**
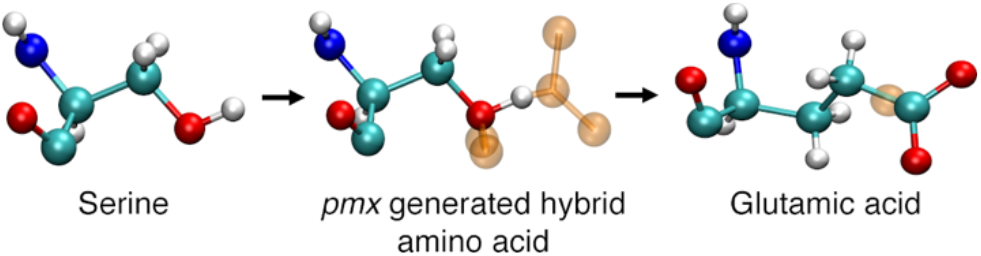
Example of a *pmx* generated hybrid amino acid structure for serine (λ = 0) to glutamic acid (λ = 1). Dummy atoms are shown as transparent orange spheres.

### Free Energy Calculation and the Thermodynamic Cycle

To estimate relative binding free energy values (ΔΔ*G*), we alchemically morphed the wild-type amino acids to their mutated forms (Figure 2). This process was replicated for both the bound and unbound states as indicated by horizontal arrows in the thermodynamic cycle shown in Figure 3. We can efficiently obtain Δ*G*_1_ and Δ*G*_3_ values with high accuracy using this approach.^46–48^ By contrast, to carry out binding/unbinding simulations (vertical arrows in Figure 3) to calculate Δ*G*_2_ and Δ*G*_4_ values would be considerably more challenging and computationally expensive.

**Figure 3.**
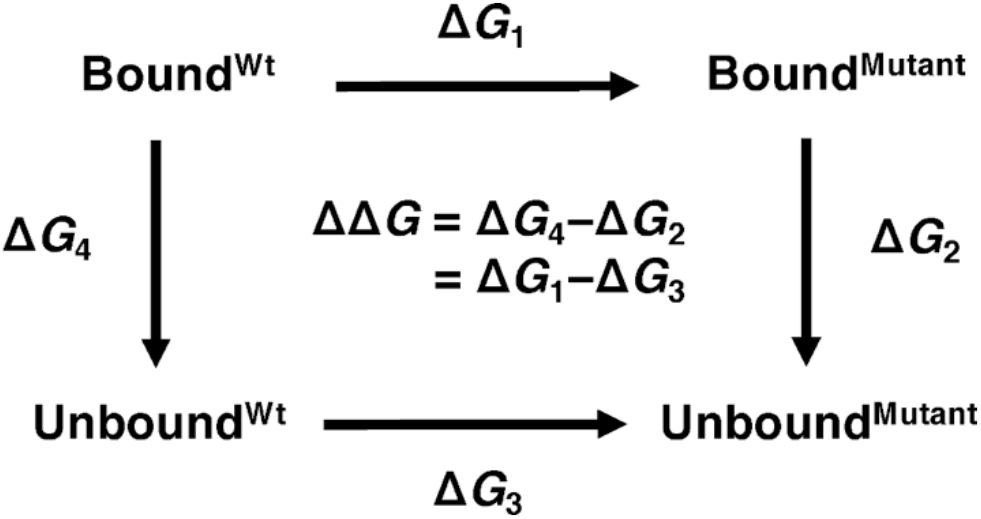
Schematic representation of the thermodynamic cycle used to calculate relative binding free energies due to mutation (ΔΔ*G*=Δ*G_1_*–Δ*G_3_*). Horizontal arrows indicate the non-physical pathways used in the current study where the amino acid was alchemically morphed from wildtype to its mutant form for both bound and unbound states.

To estimate Δ*G*_1_ and Δ*G*_3_ (two horizontal arrows in Figure 3), we used the double-system/single-box approach developed by Gapsys *et al*.^40^ Following this approach, we placed Bound^Wt^ protein complex and Unbound^Mutant^ protein in a single simulation box (λ = 0, Figure 4A) and similarly we placed Bound^Mutant^ protein complex and Unbound^Mutant^ protein in a second simulation box (λ = 1, Figure 4A). Figure 4B represents the series of steps involved for setting up the system for MD simulations and alchemical free energy calculations. The distance between the two protein systems in each simulation box was maintained at 30 Å (Figure 4B) by applying position restraints on a single backbone atom close to the center of mass of each protein system. This separation distance was chosen to be larger than the short-range electrostatics cutoff to ensure that the two protein systems in a single simulation box did not interact with each other. Alchemical transformation from λ = 0 to 1 is termed “forward”, where Bound^Wt^ was transformed into Bound^Mutant^ and simultaneously Unbound^Mutant^ was transformed into Unbound^Wt^, i.e., “backward” λ = 1 to 0. Two independent simulations (forward and backward) were thus performed to calculate ΔΔ*G* value for each mutation. Use of the double-system/single-box approach enabled us to maintain charge neutrality of the simulation system, even when an alchemical transformation involved a charge change between the wild-type and a mutant state, e.g., R83Q.

**Figure 4.**
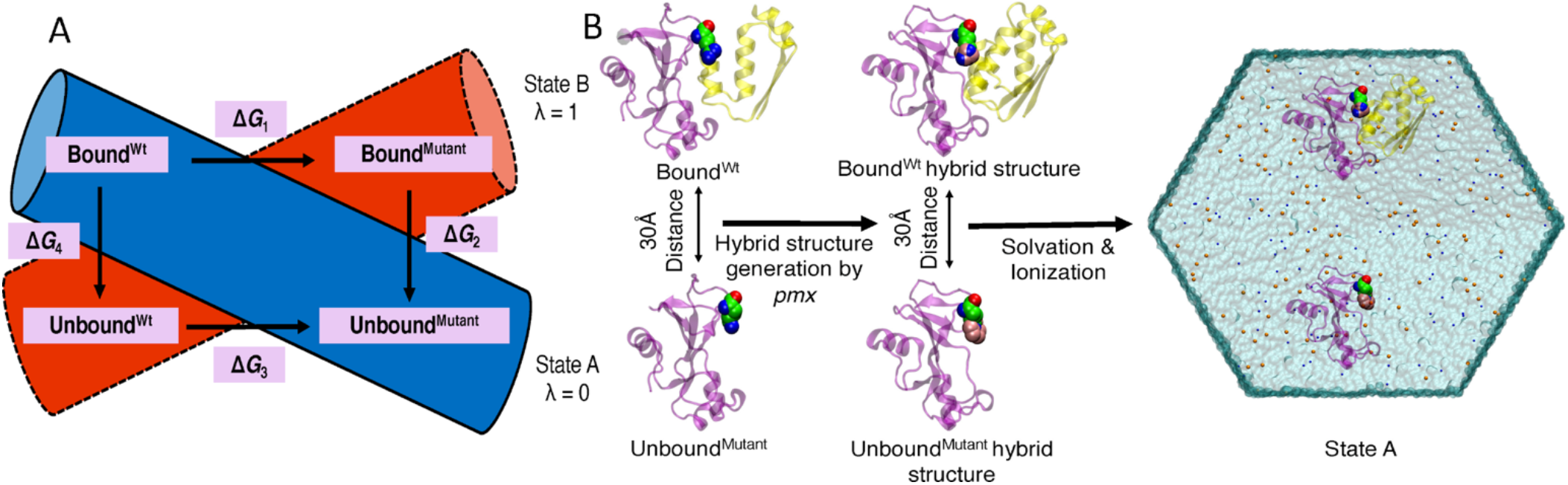
Double-system/single-box simulation setup. (A) Each colored cylinder represents a simulation box. During the forward alchemical transition, double systems consisting of Bound^Wt^ and Unbound^Mutant^ (blue cylinder, λ = 0) are morphed into Bound^Mutant^ and Unbound^Wt^ (λ = 1) states respectively. Similarly, backward alchemical transition (λ = 1 to λ = 0) takes place in the red cylinder. (B) Schematic representation of the steps involved for setting up one the double-system/single-box simulations for a mutation of 1BRS protein complex.

### MD simulations and Alchemical Free Energy Calculations

All MD simulations were carried out with the GROMACS-2018.3^49^ MD simulation package using the Amber99SB*ILDN force field, and the TIP3P water model.^50^ The *pmx*-generated hybrid structures and modified forcefield files were used as an input. For each mutation, we prepared two simulation boxes (λ = 0 and λ = 1, Figure 4A) to carry out forward and backward transitions using the steps shown in Figure 4B. Both the states were solvated using dodecahedron water boxes. Na^+^, and Cl^−^ ions were added at a 0.15 M concentration to neutralize the net charge. Both the simulation boxes were then energy minimized for 10,000 steps using the steepest descent algorithm. Subsequent NVT followed by NPT ensemble simulations were performed for 500 ps for each simulation box. During the MD simulation, constant pressure and temperature were maintained using Parrinello-Rahman^51^ pressure coupling at 1 atm and v-rescale temperature^52^ coupling at 300 K. A 2 fs time step was used and each snapshot was saved at every 10 ps. Final production MD simulations were then performed for 40 ns to ensure sufficient sampling under NPT conditions. To prevent the diffusion of the proteins and maintain a 30 Å distance between the two protein systems, backbone carbons close to the center of mass were harmonically restrained with a force constant of 1000 kJ/mol nm^2^. Choice of backbone C atoms used to apply position restraints for 1BRS was made based on the bound and unbound forms: i) site A40 of bound-state Barstar; ii) site A74 of unbound Barnase and; iii) site L20 of unbound Barstar. While for 3HFM, i) site Q37 of the bound-state light chain; ii) site H41 of unbound state of light chain iii) site L56 of the antigen. The light chain is always bound to the heavy chain regardless of whether the antigen is bound or unbound. These positional restraints affect only the translational degrees of freedom of the proteins, not the overall structure or orientation of the proteins. The contribution of the positional restraints to the estimation of Δ*G* will be the same for bound and unbound form of the proteins and thus the bias cancels out when calculating ΔΔ*G*, as is the case for the current study.

After the equilibrium MD simulations, fast-growth non-equilibrium alchemical simulations were performed to estimate the ΔΔ*G*. From each equilibrated MD simulation, the first 10 ns of the trajectory was discarded, and the last 30 ns was used to generate 100 snapshots (i.e., every 300 ps). Each snapshot was used to initialize a non-equilibrium simulation with a transition time of 5 ns where λ was continuously changed from 0 to 1 or from 1 to 0. The speed of λ value change was set 2e-7/fs for all forward and backward transitions. The derivatives of the Hamiltonian with respect to λ were recorded at every step and free energies were calculated from the work (*W*) distributions obtained from integration according to equation (1).

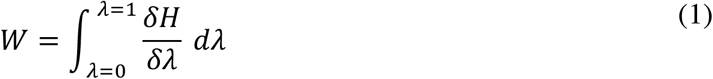

ΔΔ*G* was estimated by calculating the intersection of the forward and backward work distributions according to the Crooks-Gaussian-intersection (CGI) method as described in Goette and Grubmüller.^53^ The scripts used for analysis and calculations of ΔΔ*G* were obtained from the *pmx* package.

## RESULTS AND DISCUSSION

The purpose of our study is to test the accuracy of using *pmx* hybrid topologies and alchemical free energy calculations with the double-system/single-box approach developed by Gapsys *et al*. to estimate relative binding affinities of protein-protein complexes. The *pmx* package allows for automated generation of the necessary hybrid topologies that are otherwise challenging to generate, and the double-system/single-box approach is a simple approach to maintain a neutral charge even when a mutation changes the protein charge. We tested this approach on two proteinprotein systems of varying sizes (1BRS and 3HFM). For each system, we selected eight distinct mutations with experimental ΔΔ*G* values reported in the literature using the criteria listed under the Methods section.

For alchemical non-equilibrium free energy calculations using the fast growth method^39,54,55^ the transition time from λ = 0 to 1 or λ = 1 to 0 significantly influences the accuracy of ΔΔ*G* prediction. Short transition times lead the system far away from the equilibrium leading to a heavily biased estimate, while long transition times are less biased but more computationally costly, so the right balance is required.^39^ To develop our simulation protocol, we initially chose two mutations from the 1BRS protein complex as test cases. These cases represent the most stabilizing (D54A, ΔΔ*G* = −0.53 kcal/mol) and destabilizing (D39A, ΔΔ*G* = 6.79 kcal/mol) charge changing mutations from the list of eight selected mutations for 1BRS protein complex. (see Table 1). To determine a reasonable transition time for our production simulations, we calculated ΔΔ*G* values for both the test case mutations of 1BRS using 100 transitions with a range of transition times from 1 to 7 ns. SI Fig 1 shows that a transition time of 5 ns was sufficient to accurately estimate the free energies for these challenging mutations.

**Table 1.**
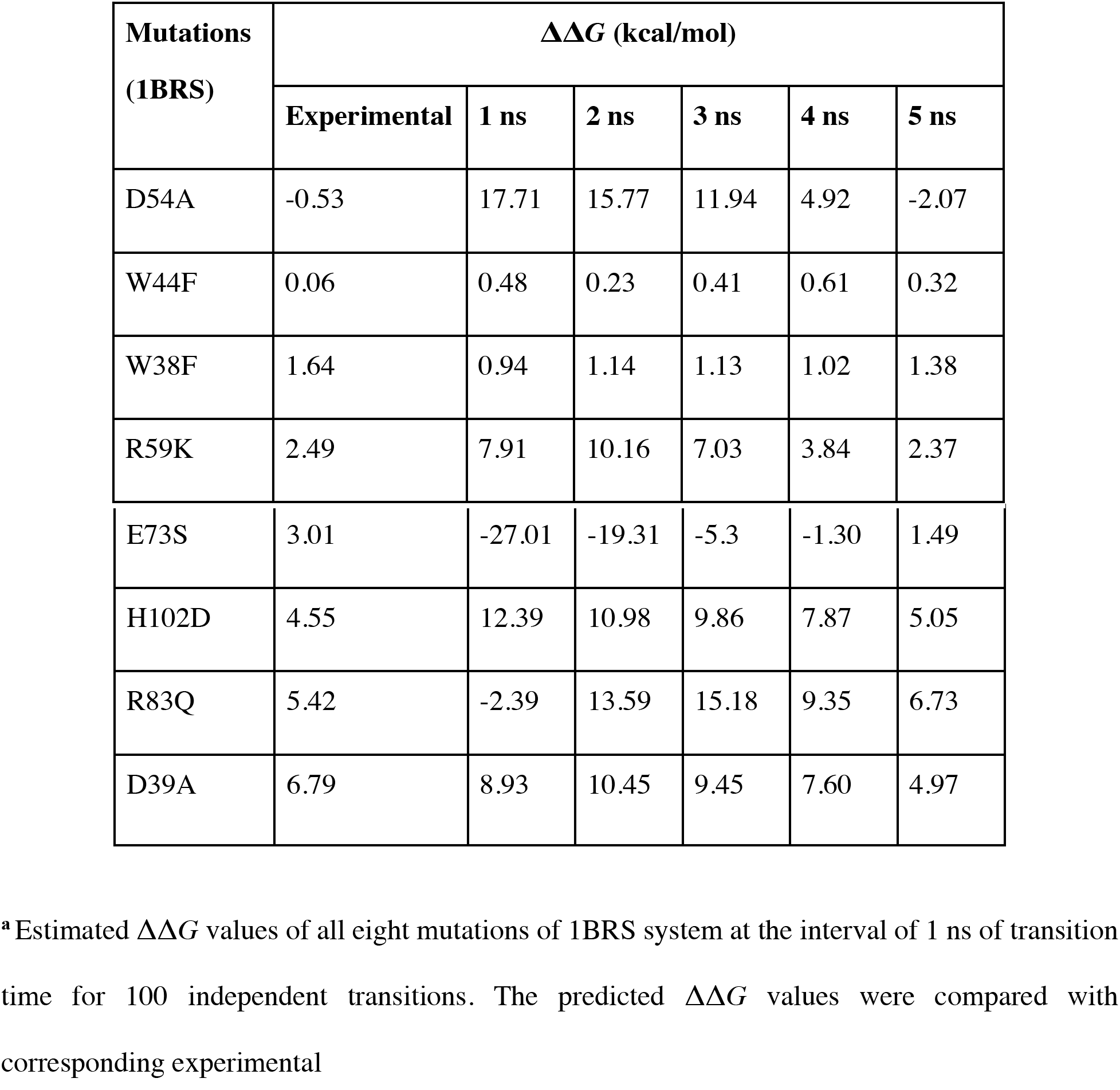
Predicted relative binding free energy of each mutation of 1BRS at different transition time between 1 to 5 ns for 100 independent transitions^a^

ΔΔ*G* values of the remaining six mutations of 1BRS were estimated using the optimized simulation protocol and the optimum transition time established through test case mutations. The predicted ΔΔ*G* values were within ± 2 kcal/mol of experimental ΔΔ*G* values for a 5 ns transition time. We used the same protocols of 40 ns of equilibration simulation with transition time of 5 ns/snapshot for 100 snapshots to predict ΔΔ*G* for eight distinct mutations of the 3HFM system using non-equilibrium fast-growth method.

Figure 5 shows the correlation between the predicted and experimental ΔΔ*G* values for all mutations from both the test systems. For 1BRS, the calculated ΔΔ*G* values correlate well with experimental data (R^2^ = 0.85). By contrast, for the antigen-antibody complex 3HFM, the correlation is significantly lower (R^2^ = 0.36). The non-charge mutations from 1BRS system such as W44F and W38F have the predicted ΔΔ*G* values within range of ± 0.5 kcal/mol of experimental Δ*G* values. The convergence time for these mutations was within 1-2 ns transition time/snapshot. In case of 3HFM, the non-charge mutations, Y20F and W98F have higher accuracy compared to other non-charge mutation, Y50L. Conversely, the charge changing mutations are challenging to achieve convergence in free energy calculations with short transition time. Longer transition times are likely needed in these cases to allow for sufficient conformational sampling. All the charge-changing mutations of 1BRS system converged at around a 5 ns transition time with relatively high accuracy (± 2 kcal/mol of experimental ΔΔ*G*). However, in 3HFM, the charge changing mutations D32N, R21A and D101K have higher accuracy compared to N31E and K97D. Thus, with 3HFM system the accuracy of the methods appears to be independent of whether a mutation involves a change in charge.

**Figure 5.**
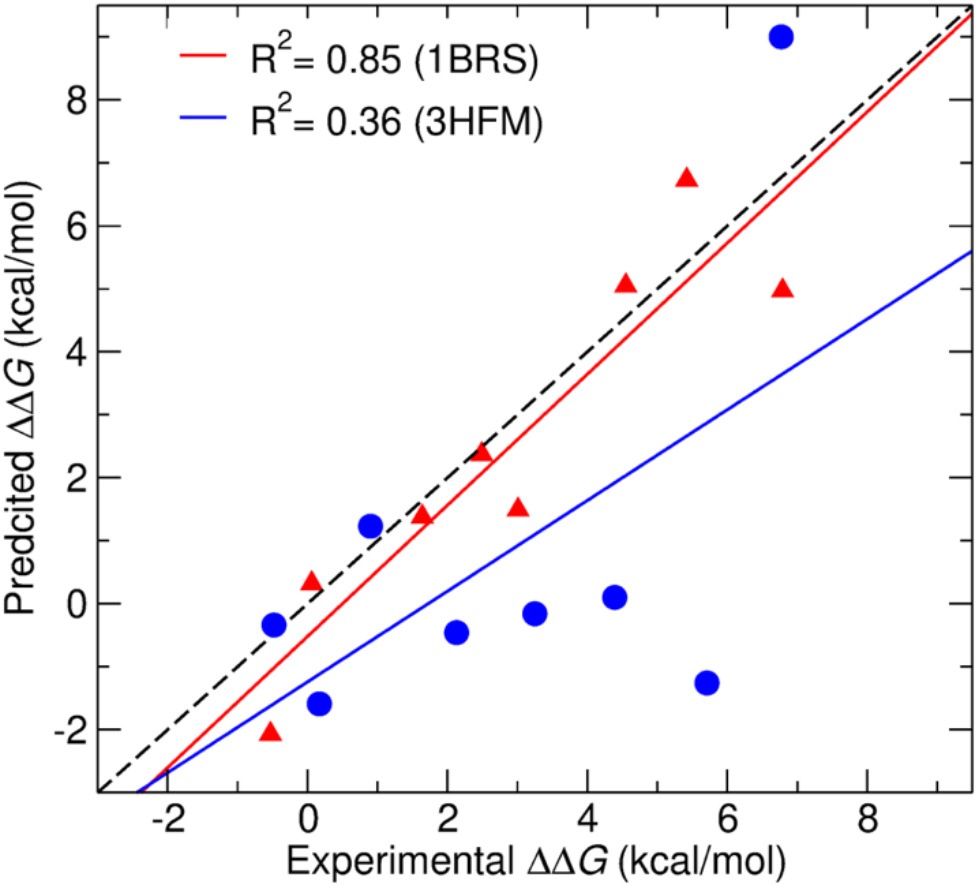
Correlation between predicted and experimental ΔΔ*G* values for 1BRS (red) and 3HFM (blue) systems. The dashed black line shows perfect correlation.

Both the test systems in this study were previously used by our lab to predict ΔΔ*G* values for the same eight mutations using the non-rigorous methods FoldX and MD+FoldX, and rigorous coarsegrained umbrella sampling MD simulations.^56^ The p*mx* with double-system/single-box approach clearly outperforms our previous FoldX^12,13^ (1BRS:R^2^=0.59, 3HFM:R^2^=-0.005) and MD+FoldX^57–59^ (1BRS:R^2^=0.62, 3HFM:R^2^=0.04) estimates in both the complexes. Interestingly, the all-atom *pmx* with double-system/single-box approach in both the complexes performs similarly to our previous coarse-grained (1BRS:R^2^=0.85, 3HFM:R^2^=0.35) method.

In this study, we used 100 snapshots per mutation to initiate the alchemical transitions and each snapshot was simulated for 5 ns. This means that 500 ns total simulation time was used to estimate Δ*G* for both forward and backward directions. The equilibration simulation required ~4500 CPUh for one mutation for 1BRS system while in case of 3HFM it required ~85,300 CPUh. With *pmx* with double-system/single-box approach, the alchemical non-equilibrium simulation time is the major contributing factor to estimate the computational cost for the calculation of one ΔΔ*G*. In 1BRS system, non-equilibrium simulations required ~45,000 CPUh for 100 transitions per ΔΔ*G* prediction, however almost 20 times more CPUh (~853,000) required in case of 3HFM system. It should also be noted that non-equilibrium alchemical transition is trivially parallelizable in that each of the 100 transitions can be run independently without relying on the completion of the previous simulation.

In order to obtain accurate binding free energy values for protein-protein complex, exhaustive conformational sampling is required in order to sufficiently explore conformational space. Larger protein-protein complexes, such as antigen-antibody complex 3HFM studied here, may require longer simulations to obtain convergence compared to smaller protein-protein complexes such as 1BRS.^60–62^ In our study, we first optimized the protocol to calculate ΔΔ*G* values for mutations of 1BRS system and then applied the same protocol to 3HFM system. It is possible that conformational sampling is one reason for obtaining lower accuracy for the 3HFM system. We note that the accuracy of the non-equilibrium method may be improved^39^ via i) longer equilibrium simulations to generate snapshots with more distant conformations, ii) increasing the transition time per snapshot, iii) increasing number of independent transitions.

Future work could involve using the alchemical double-system/single-box method but with coarse-grained protein models. Based on results from our previous study,^56^ this may significantly reduce computational cost and still retain similar accuracy. However, coarse-grained hybrid topologies of the proteins have not yet been developed. Another approach to reducing computational cost could be use of a dual resolution water model where water around the protein is atomistic and the rest of the water molecules coarse-grained.^63–65^

## CONCLUSION

In this study, we have estimated protein-protein relative binding affinities due to single amino acid mutations using *pmx* hybrid topologies with a double-system/single-box approach. Non-equilibrium alchemical methods were used to generate ΔΔ*G* estimates for one small and one large protein-protein complex, and results were compared with experimental values. We obtained a significantly higher correlation between predicted and experimental ΔΔ*G* values for the small complex compared to the larger one. We were able to distinguish stabilizing mutations from non-stabilizing mutations for all mutations in small complex, but only four out of eight for the large. The accuracy of the predictions for the large complex is lower but is similar to previously tested rigorous and non-rigorous methods. Our results suggest that there are still potential areas for improvement in the accuracy of binding free energy calculations, especially for larger proteinprotein complexes. Future work could also be devoted to estimating binding free energies due to multiple mutations.

## ASSOCIATED CONTENT

### Supporting Information

The following files are available free of charge. Prediction of ΔΔ*G* values of test mutations of 1BRS system as a function of transition time (.docx)

## AUTHOR INFORMATION

### Notes

The authors declare no competing financial interest.

