## Supplemental Figure 1 for "Implementing and assessing an alchemical method for calculating protein-protein binding free energy"

**SI Contents**

1. Prediction of ∆∆*G* values of test mutations of 1BRS system as a function of transition time.
1. **Prediction of ∆∆*G* values of test mutations of 1BRS system as a function of transition time**

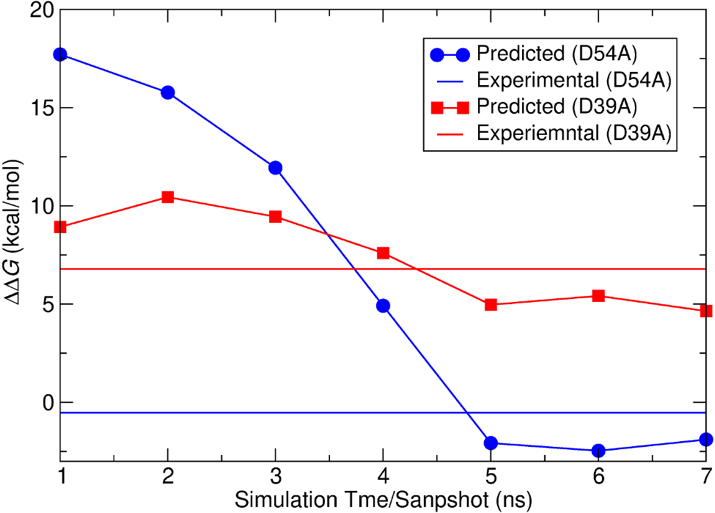

**Figure S1. Predicted ∆∆*G* values of D54A (blue) and D39A (red) mutations as a function of transition time for 100 independent transitions.** ∆∆*G* values of two test mutations D54A and D39A from 1BRS system were predicted at different transition time of 1 ns to 7 ns. The predicted ∆∆*G* values for D54A and D39A were compared with corresponding experimental ∆∆*G* values (shown in solid dashed line).
